# The upstrap

**DOI:** 10.1101/262436

**Authors:** Ciprian M. Crainiceanu

## Abstract

Bootstrap [2] is a landmark method for quantifying variability. It uses sampling with replacement with a sample size equal to that of the original data. We propose the upstrap, which samples with replacement either more or fewer samples than the original sample size. We illustrate the upstrap by solving a hard, but common, sample size calculation problem.

## 1. Algorithm

Consider a data set where the observed vector of observations at the subject level are *Y_i_*, *i* = 1,…, *n*, *θ* is a parameter of interest, and 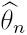 is an estimator of *θ*. For a fraction *r ∈* (0, ∞) we are interested in estimating the distribution of 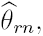 the estimator based on the fraction *r* of the original sample size. The upstrap algorithm is as follows

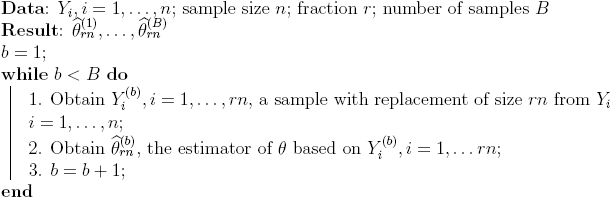

## 2 Sample size calculation for regression

We now show how to calculate the sample size using upstrapping in a relatively simple regression scenario for which there are no standard methods. Consider the case of a binary regression problem where the outcome is whether or not a person has moderate to severe sleep apnea and the predictors are gender, age, BMI, hypertension status, and hypertension by age interaction. The data comes from the Sleep Heart Health Study (SHHS) [3, 4] and is publicly available as part of the The National Sleep Research Resource (NSRR) [1]. Moderate to severe sleep apnea is defined as a Respiratory Disturbance Index at ≥4% oxygen desaturation (rdi4p) greater or equal to 15 (citations). We use data from visit 1 of the SHHS, which contains 5804 individuals.

**Figure 1:**
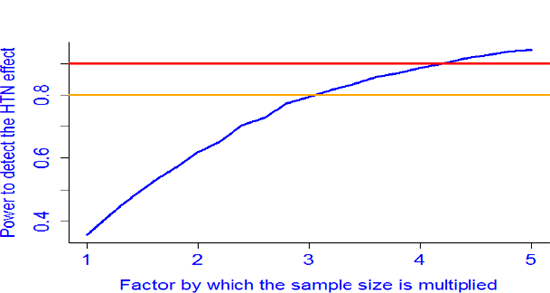
Power to detect the HTN effect in an upstrap (sample with replacement of the data with a larger sample size as the original data) as a function of the multiplier r of the original sample size.

The regression results based on these data are shown below indicating that hypertension (coded `HTNDerv_sl`) is not significant at the *α* = 0.05 level.

|  | Estimate | Std. Error | z value | Pr(>\|z\|) |  |
| --- | --- | --- | --- | --- | --- |
| (Intercept) | -8.710927 | 0.421770 | -20.653 | < 2e-16 | *** |
| gender | 1.156897 | 0.079181 | 14.611 | < 2e-16 | *** |
| age_s1 | 0.036892 | 0.005000 | 7.378 | 1.61e-13 | *** |
| bmi_s1 | 0.138369 | 0.007412 | 18.669 | < 2e-16 | *** |
| HTNDerv_s1 | 0.758377 | 0.473129 | 1.603 | 0.109 |  |
| age_s1:HTNDerv_s1 | -0.008748 | 0.007146 | -1.224 | 0.221 |  |

The question that we would like to answer is at what sample size do we expect to identify a hypertension effect on having moderate to severe sleep apnea in this model using the two-sided Wald test at *α* = 0.05 with a probability 1 – *β* = 0.8. The idea is simple. We set a grid of fractions of the sample size; in this case this grid is *r* = 1,1.2,…, 5 and for every value of r we upstrap *B* = 10000 data sets of size *rn.* For every sample we conduct the two-sided Wald test for HTN in the model above and reject the null hypothesis of no association if the corresponding *p*-value is less than 0.05. Figure 1 provides the frequency with which the test for no HTN effect is rejected as a function of the multiplier of the sample size, *r*. For example, for multiplier *r* = 2 we produced *B* = 10000 samples with replacement from SHHS data sets with twice the number of subjects 2*n* = 11608. For each data we ran the model and recorded whether the *p*-value for HTN was smaller than 0.05. For this sample size we obtained that the HTN effect was identified in 62% of the samples. We also obtained that the power was equal to 0.794 at the sample size mutiplier *r* = 3.0 and 0.817 at multiplier *r* = 3.2, indicating that the power 0.8 would be attained at 3.1 * *n* ≈ 17,500 subjects. There are very few methods to estimate the sample size in such examples and we contend that the upstrap is a powerful and general method to conduct such calculations. Similar approaches could be used in many other situations, including estimating a fixed effect (e.g. treatment) using longitudinal data in the context of a clinical trials or the sample size necessary to detect gene by gene and gene by environment interactions in genomics studies.

